# Anticodon Table of the Chloroplast Genome and Identification of Putative Quadruplet Anticodons in Chloroplast tRNAs

**DOI:** 10.1101/2022.04.24.489293

**Authors:** Tapan Kumar Mohanta, Yugal Kishore Mohanta, Ahmed Al-Harrasi, Nanaocha Sharma

**Affiliations:** Natural and Medical Sciences Research Center, University of Nizwa, Nizwa, 616, Oman; Dept. of Applied Biology, School of Biological Sciences, University of Science and Technology Meghalaya, Baridua, 793101, Meghalaya, India; Institute of Bioresources and Sustainable Development, 795001, Imphal, Manipur, India

**Keywords:** Chloroplast, tRNA, Anticodons, Evolution, Quadruplet anticodons

## Abstract

The chloroplast genome of 5959 species was analyzed to construct the anticodon table of the chloroplast genome. Analysis of the chloroplast transfer ribonucleic acid (tRNA) revealed the presence of a putative quadruplet anticodon containing tRNAs in the chloroplast genome. The tRNAs with putative quadruplet anticodons were UAUG, UGGG, AUAA, GCUA, and GUUA, where the GUUA anticodon putatively encoded tRNA^Asn^. The study also revealed the complete absence of tRNA genes containing ACU, CUG, GCG, CUC, CCC, and CGG anticodons in the chloroplast genome from the species studied so far. The chloroplast genome was also found to encode tRNAs encoding N-formylmethionine (fMet), Ile2, selenocysteine, and pyrrolysine. The chloroplast genomes of mycoparasitic and heterotrophic plants have had heavy losses of tRNA genes. Furthermore, the chloroplast genome was also found to encode putative spacer tRNA, tRNA fragments (tRFs), tRNA-derived, stress-induced RNA (tiRNAs), and group I introns. An evolutionary analysis revealed that chloroplast tRNAs had evolved via multiple common ancestors and the GC% had more influence toward encoding the tRNA number in the chloroplast genome compared to the genome size.

## Introduction

The origin of the genetic code and the translation event are considered to be major transition points in the evolution of biology. The triplet genetic code is hailed as one of the most important and ultimate evolutionary anchors and an indisputable piece of evidence of life. The triplet genetic code understands the specific assignment of the amino acids in the translation machinery. It is the universal manual and a guided dictionary that cells use to translate the corresponding amino acids into the translating protein. The number of codon combinations on the mRNA can be an astounding number of feasible protein sequences, from which only a few can be found in nature. It is believed that the triplet genetic code is universal and degenerate, and accommodates twenty essential amino acids using sixty-one sense and three stop codons. However, an emerging study has proved that the “Universal Genetic Code” is no more universal and can be called as canonical [1, 2]. Sometimes nature enhances the protein functionalities through codon reassignment to incorporate new amino acids. This has led to the discovery of the role of selenocysteine (Sec) and pyrrolysine (Pyl) amino acid in the protein, through the assignment of the stop codon as the sense codon. However, sense codon reassignment requires low frequency codons, and hence, stop codons are used for this purpose. It has been demonstrated that except for the triple codons, the Escherichia coli ribosome can accommodate codons and anticodons of variable sizes [3]. Taking this opportunity they have translated four base codon pairs CCCU, AGGA, UAGA, and CUAG using the four base anticodons [3]. Frame-shifting of the +1 nucleotide is most favorable in the absence of suppressor tRNA in E. coli [3]. The study also reveals that frame maintenance during translation is not absolute and a frame shift can be promoted by mutant tRNAs and it can occur with high frequencies at the programmed site of the mRNA [4, 5]. Riddle and Carbon (1973) reported the presence of four base anticodons CCCC in tRNA^Gly^ instead of the wild-type CCC [6]. A study conducted by Mohanta et al., (2020) revealed the presence of nine nucleotide anticodons instead of seven nucleotides [7]. These features in the tRNAs certainly explain the presence of extended codons and anticodons. Most possibly these kind of evolutionary scenarios exist in codons and tRNAs, to meet the novel translational demand.

The availability of enormous genome sequencing data is quite valuable to dig deep into the molecular features of the protein translation machinery. Significant studies have been performed in the field of codons and tRNAs (anticodons) and yet, a number of things need to be explored. Taking this opportunity, we have conducted a large-scale study to deduce the anticodon table of the chloroplast genome, to understand the presence of reduced or extended genetic codes/anticodons in tRNAs. Furthermore, we have also tried to understand the presence of Sec and Pyl tRNAs, which are part of the extended genetic code. A Furthermore investigation has also been conducted to understand the presence of different introns and the presence of a possible spacer tRNA and tRNA fragments.

## Results

### tRNAs with ACU, CUG, GCG, CUC, CCC, and CGG anticodons are absent in the chloroplast genome

Analysis of the chloroplast genome of the 5959 species from Algae (303), Bryophyte (69), Eudicot (3832), Gymnosperm (153), Magnoliids (182), Monocot (1177), Nymphaeales (34), protist (57), Pteridophyte (139), and unknown (13) led to the discovery of 215966 tRNA genes. We did not find any tRNA encode for ACU, CUG, GCG, CUC, CCC, and CGG anticodons from them (Table 1). Furthermore, we also found several anticodons, which were seemingly very rare in the chloroplast genome. They were AGU (tRNA^Thr^), AAG (tRNA^Leu^), CGC (tRNA^Ala^), UCA (tRNA^Sup^), AGG (tRNA^Pro^), AUU (tRNA^Asn^), UAU (tRNA^Ile^), AUA (tRNA^Tyr^), CAG (tRNA^Leu^), CUU (tRNA^Lys^), CCU (tRNA^Arg^), AAU (tRNA^Ile^), and GAG (tRNA^Leu^) (Table 1, Supplementary File 1). The tRNA with anticodons AAG, AGU, and CGC was found only once, whereas, the tRNA with anticodon UCA, AGG, and AUU was found twice for each (Table 1, Supplementary File 1, Supplementary File 2). However, the percentage of the CAU (5.47%, tRNA^Met^) anticodon was the highest among all the 64 anticodons. The abundance of the CAU anticodon was followed by GUU, UGC, ACG, and others (Table 1).

**Table 1.** Anticodon Table of the chloroplast genome. Study from 5959 chloroplast genomes shows several rare anticodons.

|  |  |  |  |  |  |  |
| --- | --- | --- | --- | --- | --- | --- |
| Ala | AGC: 214 | GGC: 203 | CGC: 1 | UGC: 11017 |  |  |
| Arg | ACG: 10927 | GCG: 0 | CCG: 304 | UCG: 103 | CCU: 12 | UCU: 5729 |
| Asn | AUU: 2 | GUU: 11018 |  |  |  |  |
| Asp | AUC: 399 | GUC: 6122 |  |  |  |  |
| Cys | ACA: 224 | GCA: 6156 |  |  |  |  |
| Gln | CUG: 0 | UUG: 6242 |  |  |  |  |
| Glu | CUC: 0 | UUC: 5925 |  |  |  |  |
| Gly | ACC: 204 | GCC: 4917 | CCC: 0 | UCC: 5684 |  |  |
| His | AUG: 405 | GUG: 7148 |  |  |  |  |
| Ile | AAU: 22 | GAU: 10695 | CAUIle2: 10575 | UAU: 5 |  |  |
| Leu | AAG: 1 | GAG: 42 | CAG: 6 | UAG: 5745 | CAA: 10686 | UAA: 5546 |
| Lys | CUU: 7 | UUU: 5312 |  |  |  |  |
| Met | CAU Met: 11741<br>CAUfMet: 709 |  |  |  |  |  |
| Phe | AAA: 207 | GAA: 5982 |  |  |  |  |
| Pro | AGG: 2 | GGG: 817 | CGG: 0 | UGG: 5883 |  |  |
| Ser | AGA: 203 | GGA: 5520 | CGA: 173 | UGA: 5174 | ACU: 0 | GCU: 5980 |
| Thr | AGU: 1 | GGU: 5703 | CGU: 454 | UGU: 5669 |  |  |
| Trp | CCA: 5955 |  |  |  |  |  |
| Tyr | AUA: 6 | GUA: 5964 |  |  |  |  |
| Val | AAC: 399 | GAC: 10687 | CAC: 200 | UAC: 4748 |  |  |
| SeC | UCA: 204 |  |  |  |  |  |
| Pyl | CUA: 197 |  |  |  |  |  |
| Sup | CUA: | UUA: 205 | UCA: 2 |  |  |  |

### Chloroplast genome encodes tRNA for N-formylmethionine, Ile2, Selenocysteine, and Pyrrolysine

A study revealed, a chloroplast genome was found to encode tRNAs for tRNA^fMet^, tRNA^Ile2^, tRNA^Sel^, and tRNA^Pyl^ (Table 1). The tRNA^fMet^ was encoded by the same CAU anticodon that coded tRNA^Met^. We found 709 (0.33%) genes that encoded tRNA^fMet^ (Table 1). Also, tRNA^Ile^ encoded by the CAU anticodon was commonly referred to as tRNA^Ile2^ (Table 1). We found at least 10575 (4.93%) tRNA genes encoding tRNA^Ile2^ (Table 1). Selenocysteine amino acid was encoded by a previously known stop codon UCA. At least, 204 chloroplast genes were found to encode the UCA anticodon for tRNA^Sel^ (Table 1, Supplementary File 3). A chloroplast genome was also found to encode 197 genes for CUA anticodons that encoded tRNA^Pyl^ (Table 1). However, we did not find any CUA anticodon that encoded the suppressor tRNA (Table 1).

### Chloroplast genome encodes putative duplet and quadruplet anticodons

We have already mentioned that the triplet genetic code is not universal, it is canonical. Therefore, it is possible that the genome might have suppressed or extended the genetic code, which is yet to be elucidated, to a greater extent. In our study, we have found that the chloroplast genome encodes the putative duplet and quadruplet anticodons (Supplementary File 4, Supplementary File 5). The annotation of tRNA with quadruplet anticodon had been found when chloroplast genomes were annotated in the GeSeq chlorobox (https://chlorobox.mpimp-golm.mpg.de/geseq.html). However, re-analysis of the tRNA with the quadruplet anticodon in tRNAscan-SE did not result in a tRNA with a quadruplet anticodon, which might be due to the default setting for identification of a tRNA with a triplet anticodon. We are the first to report the presence of duplet and quadruplet anticodons in the chloroplast genome of the plant kingdom. We found that at least 91 species were encoded quadruplet anticodons (Supplementary File 4). The quadruplet anticodons were UAUG, UGGG, AUAA, GCUA, and GUUA (Supplementary File 4). The quadruplet anticodon GUUA found in Gossypium sturtianum (NC_023218.1) putatively encoded tRNA^Asn^. Similarly, at least 13 species were found to encode duplet (two nucleotides) anticodons in the tRNAs of the chloroplast genome (Supplementary File 5). Among them, there were at least eight putative unique duplet anticodons namely UG, AG, AU, CA, GA, GG, GU, and UA (Supplementary File 5). The putative duplet anticodons might have been caused by the loss of a nucleotide from the anticodon, because, if there were duplet anticodons, the genome could encode only 16 anticodons in its genome and would not be able to accommodate all the 20 coding amino acids in the protein. However, there is a high possibility of having quadruplet anticodons in the tRNAs, because, in a quadruplet anticodon table, there are 256 possibilities to encode different amino acids into the protein (Supplementary Table 1).

### Parasitic organisms have lost the tRNA genes in their chloroplast genome

We found that some of the chloroplast genomes had lost the tRNA genes. The species that have been found to have lost the tRNA genes are Pilostyles aethiopica (NC_029235.1) (Figure 1) and Pilostyles hamiltonii (NC_029236.1) (Supplementary File 6). Pilostyles aethiopica and Pilostyles hamiltonii are endoparasitic plants. Furthermore, some other plants have encoded a fewer number of tRNAs in their chloroplast genome (Supplementary File 6). They are Asarum minus (5), Gastrodia elata (5), Sciaphila densiflora (6), Epirixanthes elongata (8), Burmannia oblonga (8), Lecanorchis japonica (8), Lecanorchis kiusiana (9), and Selaginella tamariscina (9)(Supplementary File 6). All of the mentioned species encoded less than 10 tRNA genes in their chloroplast genome. Gastrodia elata is a saprophyte, whereas, Sciaphila densiflora, Epirixanthes elongate, Burmannia oblonga, Lecanorchis japonica, and Licanorchis kiusiana are mycoheterotrophs, and Cystopteris chinensis is an endangered species.

**Figure 1.**
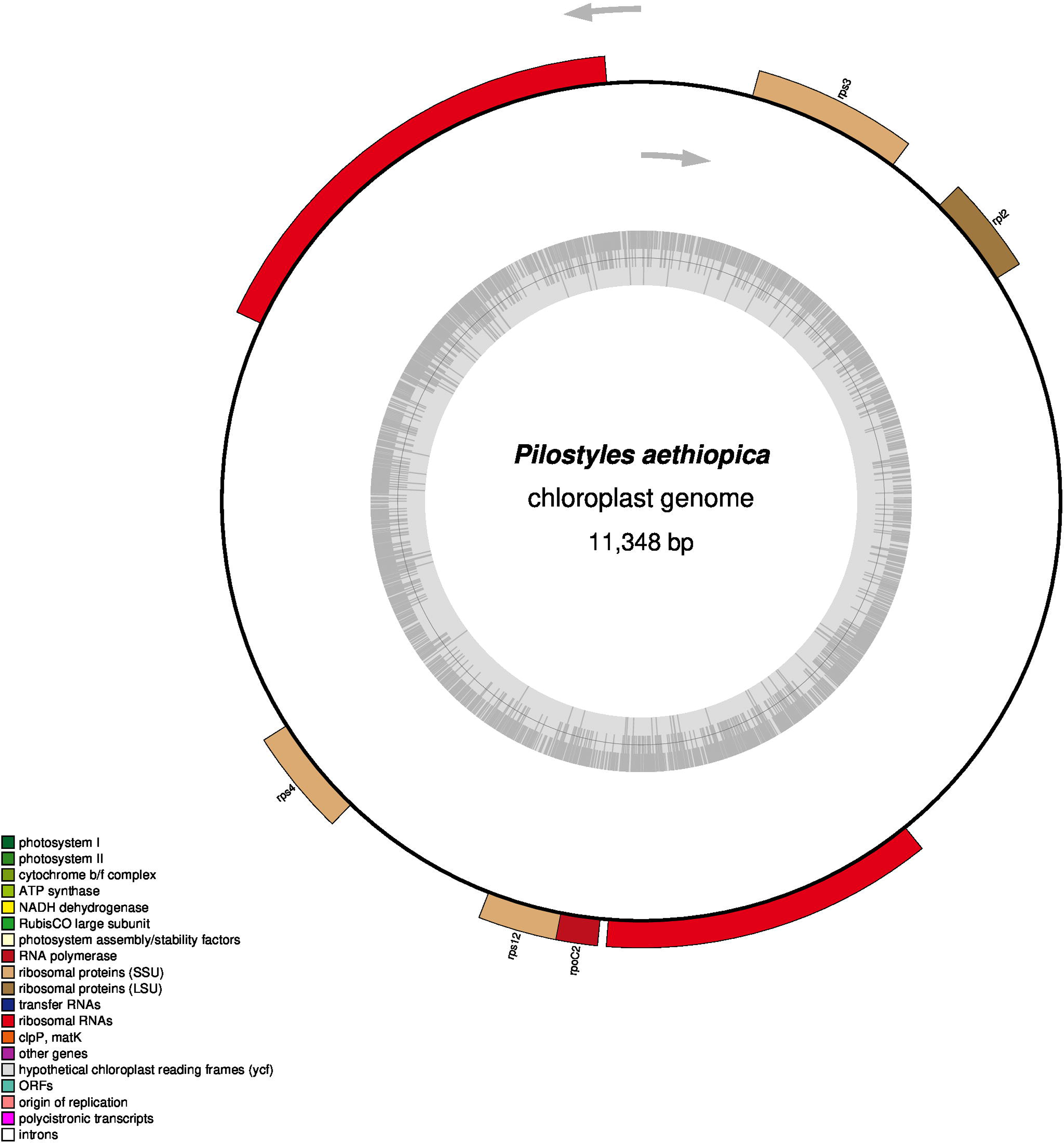
OGDRAW map of Pilostyles aethiopica (NC_029235.1) chloroplast genome. The map shows the loss of tRNA genes and inverted repeats.

The chloroplast genome of Asarum minus encoded UUU (tRNA^Lys^), UUG (tRNA^Gln^), GCU (Trna^Ser^), UCC (tRNA^Gly^), and UCU (tRNA^Arg^); Gastrodia elata encoded UUG (tRNA^Gln^), GCA (tRNA^Cys^), UUC(tRNA^Glu^), CAU(tRNA^fMet^), and CCA(tRNA^Trp^); Sciaphila densiflora encoded UUG (tRNA^Gln^), CAU (tRNA^Ile^), CCA(tRNA^Trp^), CAU(Trna^fMet^), UUC(tRNA^Glu^), and GCA(tRNA^Cys^); Epirixanthes elongata encoded CCA (tRNA^Trp^), CAU(tRNA^fMet^), UUG(tRNA^Gln^), GUC(tRNA^Asp^), GUA(tRNA^Tyr^), and UUC(tRNA^Glu^); Burmannia oblonga encoded UUG (tRNA^Gln^), GCA (tRNA^Cys^), GUA (tRNA^Tyr^), UCC (tRNA^Glu^), CAU (tRNA^fMet^), GUG (tRNA^His^), and CAU (tRNA^Ile^); Lecanorchis japonica encoded UUG (tRNA^Gln^), GCA (tRNA^Cys^), GUC (tRNA^Asp^), CAU (tRNA^fMet^), GAA (tRNA^Phe^), CAU (tRNA^Ile^), and GUU (tRNA^Asn^); Lecanorchis kiusiana encoded UUG (tRNA^Gln^), GCA (tRNA^Cys^), GUC (tRNA^Asp^), UUC (tRNA^Glu^), CAU (tRNA^fMet^), GAA (tRNA^Phe^), CAU (tRNA^Ile^), and GUU (tRNA^Asn^); and Selaginella tamariscina encoded GUG (tRNA^His^), GUC (tRNA^Asp^), GUA (tRNA^Tyr^), UUC (tRNA^Glu^), GUU (tRNA^Asn^), and CCA (tRNA^Trp^). These species encoded only 14 anticodons CAU, CCA, GAA, GCA, GCU, GUA, GUC, GUG, GUU UCC, UCU, UUC, UUG, and UUU.

### Chloroplast genome encode putative spacer tRNAs

Spacer RNA genes are usually found in the spacer region, between the 16S and 23S rRNAs, in bacterial genomes. When we focused our study on the spacer RNA in the chloroplast genome, we found that chloroplast genomes were also encoded in the putative spacer tRNAs between the 16S and 23S rRNA genes. tRNA^Ala^(UGC) and tRNA^Ile^ (GAU) were the most predominant spacer tRNAs found in the chloroplast genome (Figure 2). The percentages of the UCG and GAU anticodons in the chloroplast genome were 5.13 and 4.98, respectively. This showed that spacer tRNAs were more common in the chloroplast genome. Sometimes, it contained tRNA^fMet^ (CAU) and tRNA^Ser^ (GCU) in the spacer region. All the chloroplast genomes did not encode the spacer tRNAs (Supplementary File 7). None of a mycoparasitic plants was found to encode the putative spacer tRNA in their chloroplast genome. However, the majority of the species encoded putative spacer tRNAs.

**Figure 2.**
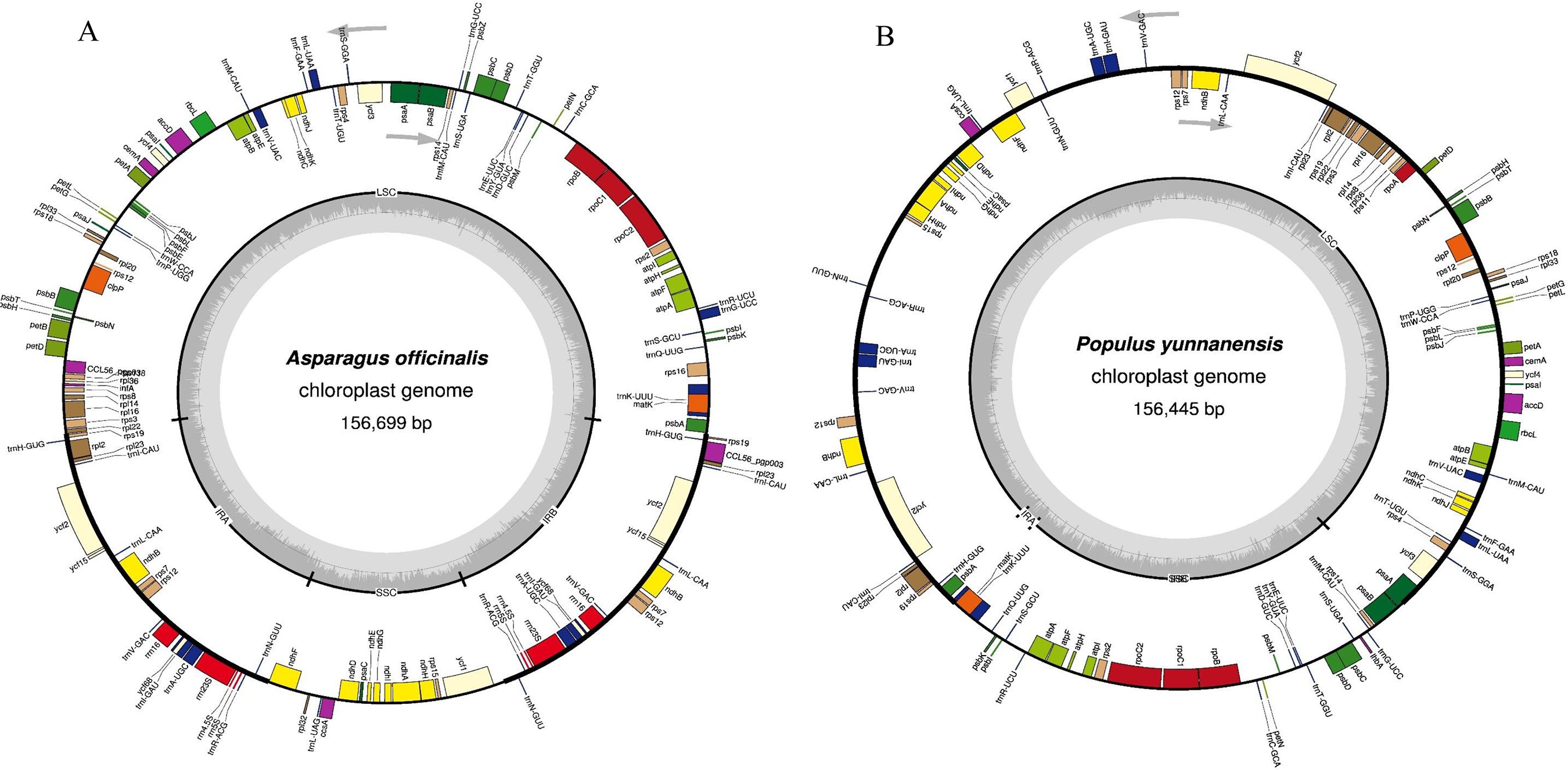
OGDRAWM map of (A) Asparagus officinalis (NC_034777.1) and (B) Populus yunnanensis (NC_037421.1) chloroplast genomes. (A) Asparagus officinalis shows the presence of a putative spacer tRNAs. tRNA^Ala^ (UGC) and tRNA^Ile^ (GAU) are present between 16S and 23S rRNA in A. officinalis. No putative spacer tRNA is found in the chloroplast genome of Populus yunnanensis.

### The Majority of chloroplast tRNAs encode group I intron

It was found that the majority of chloroplast-encoding tRNAs encode introns. Except for tRNA^Arg^, tRNA^Asn^, tRNA^Asp^, tRNA^Gln^, tRNA^His^, tRNA^Pro^, tRNA^Trp^, and tRNA^Val^ all other tRNA genes were found to contain group I introns (Table 2). The introns found in tRNA seem to be isotype-specific (Table 2). The introns are conserved within the tRNA isotype and the conserved nucleotide sequences of the introns of one isotype do not match with the conserved introns of other isotypes (Table 2). When we cluster the conserved region of the introns, they form four groups (Supplementary Figure 1). We have named them group A, B, C, and D. Group A contains tRNA^Leu^, tRNA^Tyr^, and tRNA^Cys^; group B contains tRNA^Ser^; group C contains tRNA^Lys^, tRNA^Met^, and tRNA^Ala^, and group D contains tRNA^Gly^, tRNA^Ile^, tRNA^Glu^, and tRNA^Thr^ (Supplementary Figure 1). However, the introns of tRNA^Phe^ do not group with any other introns (Supplementary Figure 1).

**Table 2.**
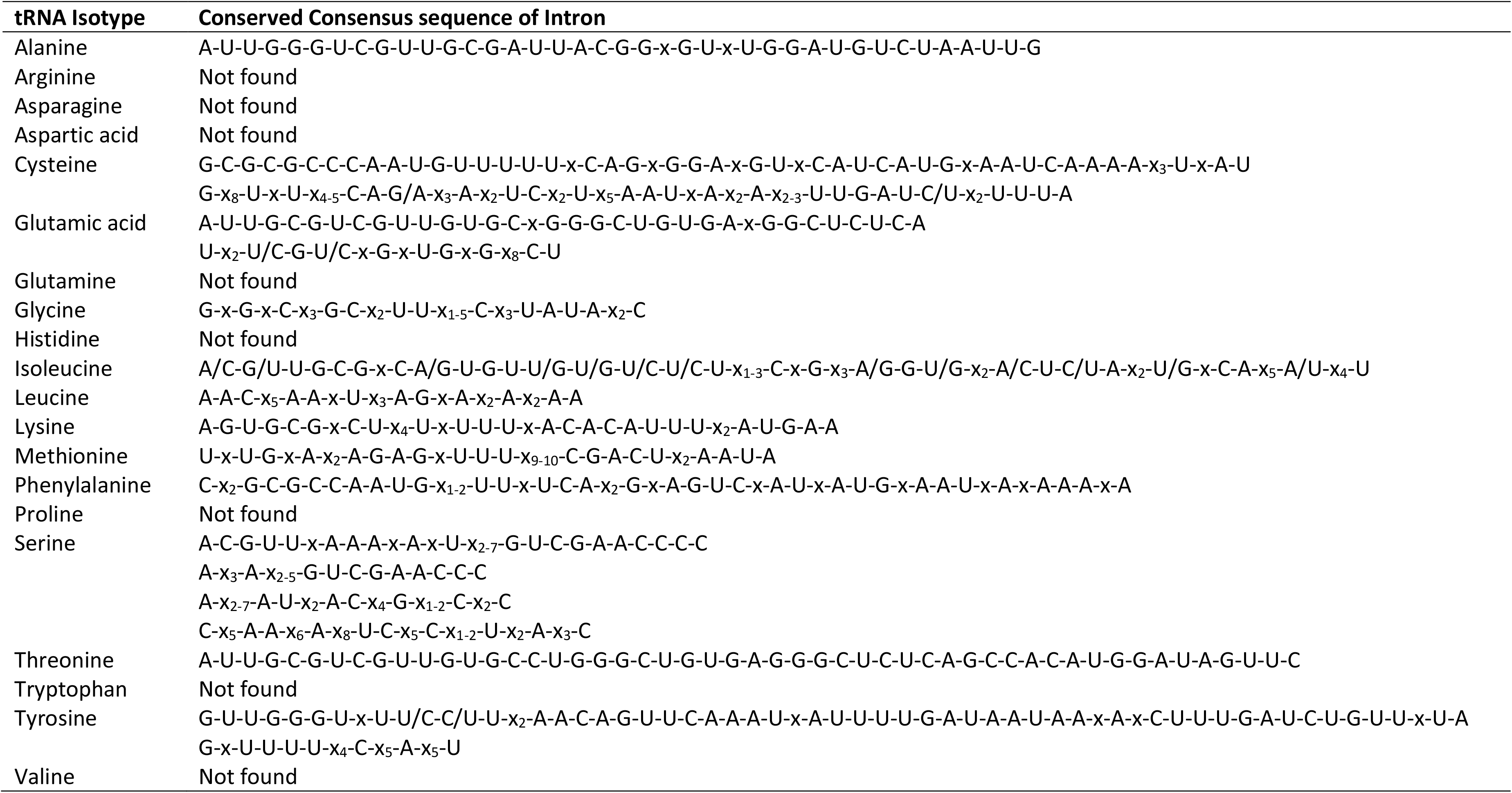
Conservation of introns in chloroplast tRNAs. From the mentioned tRNA isotypes, at least eight isotypes do not encode any intron in their tRNA genes or its not conserved.

### Chloroplast genome encode putative novel tRNAs

Although we all are well-acquainted with the fact that tRNA makes a clover leaf-like structure, yet we found some variations in the tRNA structure. Analysis revealed the presence of a few novel tRNA structure/tRNA-like molecules (Figure 3 and Figure 4). Some putative novel tRNA-like structures seemed to lack the anticodon loop, whereas, in some cases they had extra sequences near the anticodon arm region (Figure 3). A tRNA-like structure contained an extended nucleotide sequence in the region between the D-arm and anticodon arm (Figure 4). At least 42 species were found to encode novel tRNA-like structures that contained extended nucleotide sequences between the D-arm and anticodon arm (Figure 4). Furthermore, a few tRNAs were found to have lost the pseudouridine loop (Figure 5), suggesting the presence of novel tRNAs/tRNA-like structures in the chloroplast genome.

**Figure 3.**
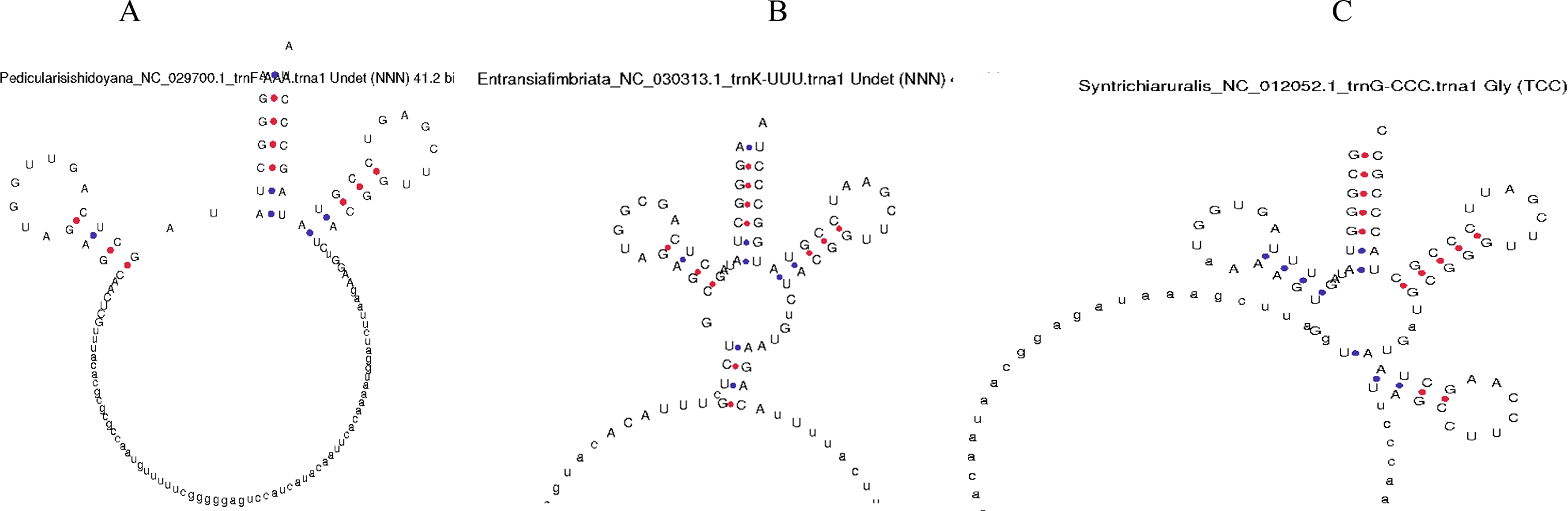
Putative novel tRNAs in chloroplast genome. (A) In the tRNA of Pedicularis ishidoyana (NC_029700.1) there is a long nucleotide sequence present in between the D arm and anticodon arm. (B) In Entransia fimbriata (NC_030313.1) tRNA (tRNA^Lys^UUU) a long nucleotide sequence is present in the anticodon loop region that masks the anticodon loop. (C) In Syntrichia ruralis (NC_012052.1) tRNA^Gly^UCC, a long nucleotide sequence is found in between the D-arm and anticodon arm.

**Figure 4.**
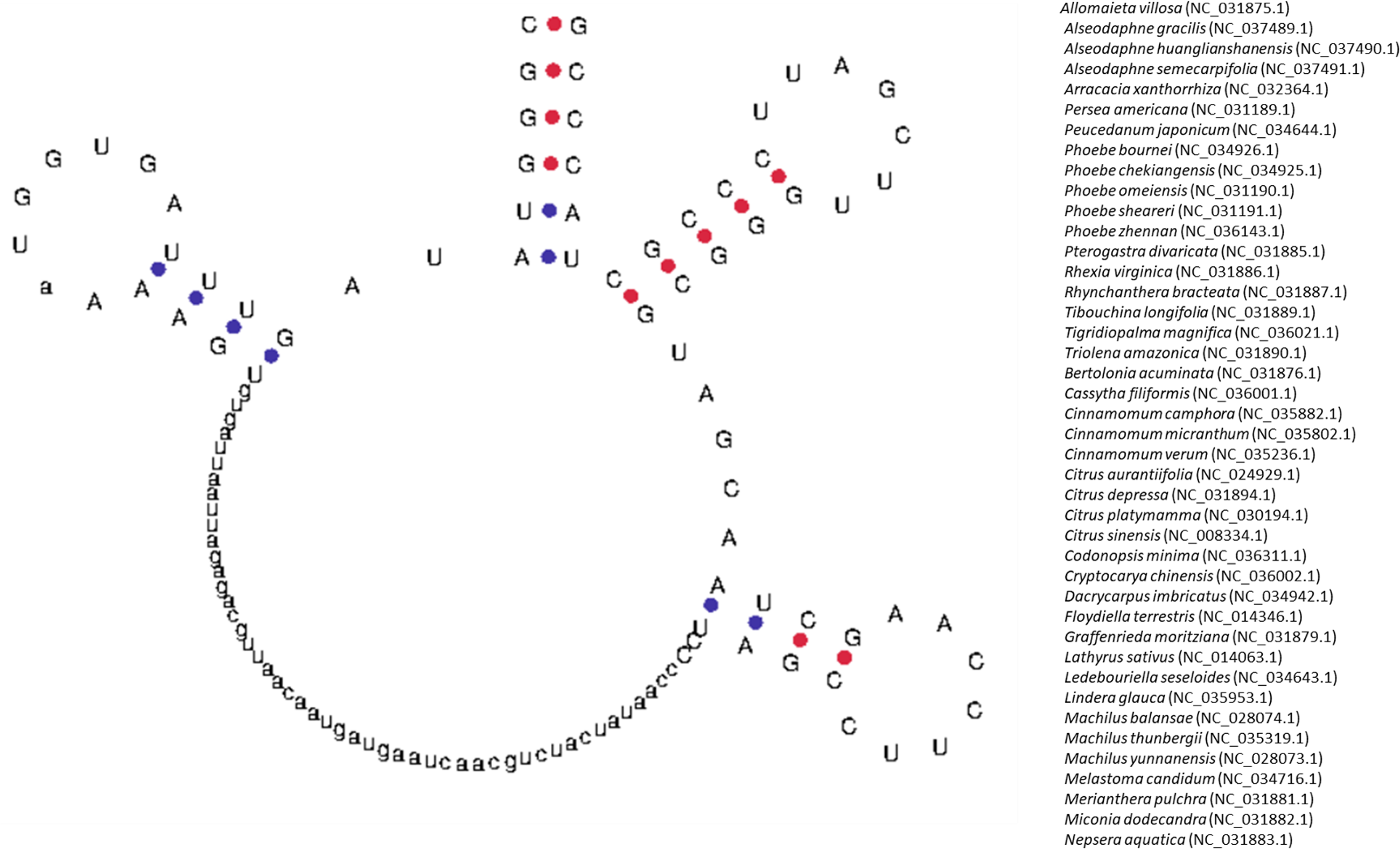
Putative novel tRNA of chloroplast tRNA. The tRNA contains a long nucleotide sequence in between the D-arm and anticodon arm. At least 42 chloroplast genomes are found to encode a similar tRNA structure in it. The structure was predicted using the tRNAscan-SE 2.0 program.

**Figure 5.**
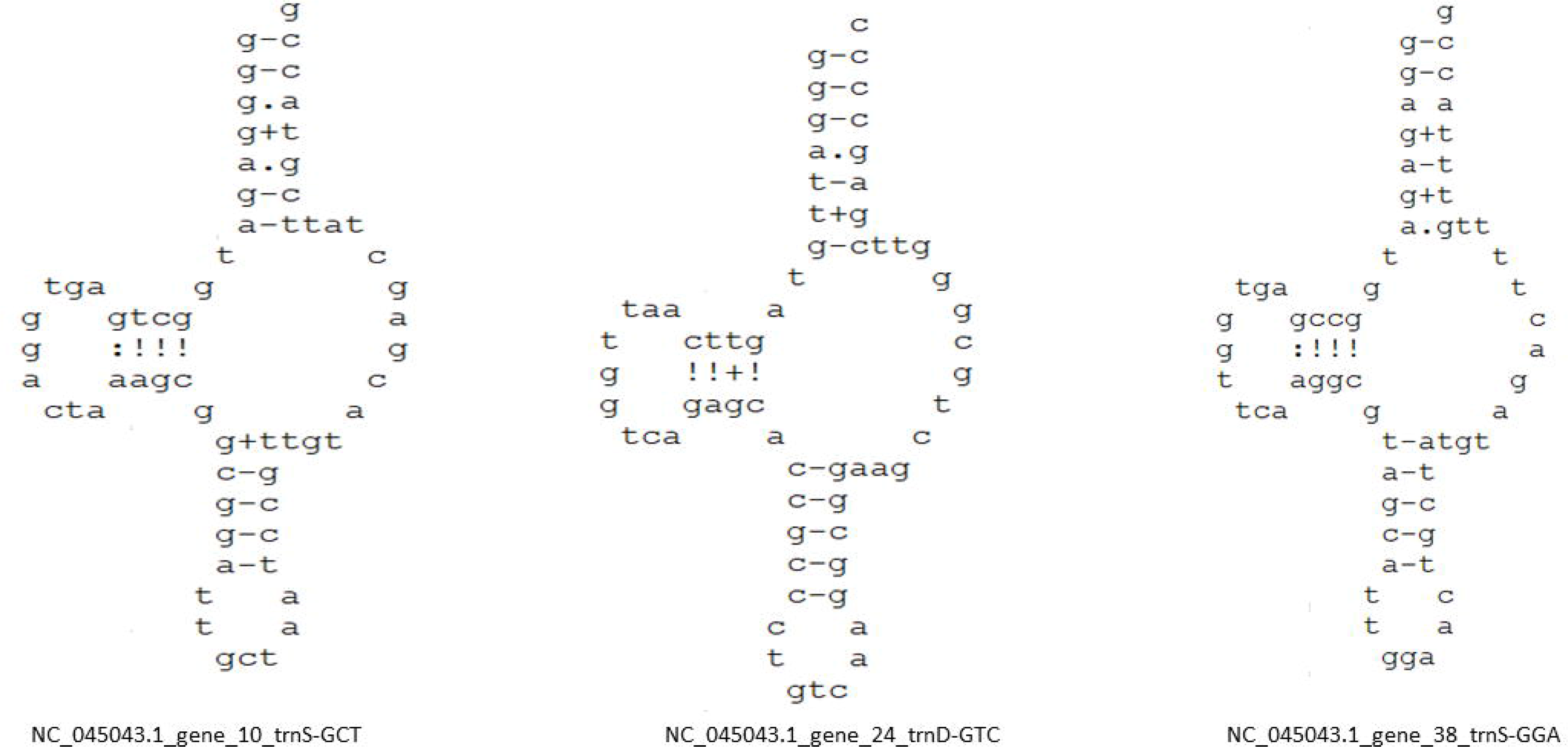
Figure 5 shows the presence of putative nematode mitochondrial tRNA in the chloroplast genome. The tRNAs have been seen to lose the Ψ-arm and Ψ-loop. The presence of nematode mitochondrial genome in the chloroplast genome shows that the truncated tRNAs are shared in between the chloroplast and mitochondria. The structure has been predicted using the Aragorn software.

### Chloroplast genome encodes putative tRNA Fragments (tRFs)

The tRFs are small 14-32 nucleotides novel class of small, non-coding RNAs, derived from the mature or precursor tRNAs that are different from the tRNA-derived, stress-induced tRNAs (tiRNAs) [8, 9]. Analysis revealed the presence of at least 55 tRFs in the chloroplast genome. The tRFs found were for tRNA^Glu^, tRNA^Arg^, tRNA^Gly^, tRNA^His^, tRNA^Val^, tRNA^Ile^, tRNA^Thr^, tRNA^Leu^, tRNA^Lys^, and tRNA^Ala^ (Supplementary File 8). The tRFs of tRNA^Glu^ were found to contain conserved nucleotide sequence GGCCTTATCGTCTAGTGAT, whereas, those of tRNA^Gly^ were found to contain conserved GCGGGTATAGTTTAGTGGTAAA nucleotides (Supplementary File 8). As such, we did not find conserved nucleotide sequences for the other tRFs. The tRFs of tRNA^Ala^, tRNA^Gly^, tRNA^Ile^, tRNA^Lys^, and tRNA^Leu^ were 5ˈ-tRFs, whereas, the tRFs of tRNA^His^, tRNA^Thr^, and tRNA^Val^ were 3ˈ-tRFs. The tRFs of tRNA^Glu^ did not match either the 5ˈ- or 3ˈ-end of the tRNA, and hence, might have originated from the precursor tRNA transcript. Therefore, they can be classified as tRF-1.

### Chloroplast genome encode putative tiRNAs

The longer tRFs (tRNA fragments) of 30–50 nucleotide-long sequences are called tRNA-derived, stress-induced RNAs (tiRNAs)[8]. Therefore, we searched for the presence of 30–50 nucleotide tRFs. We found at least 244 tRNA sequences, which encoded the 30–50 nucleotides (Supplementary File 9). The tiRFs were part of putative tRNA^Ala^ (UGC), tRNA^Phe^ (GAA), tRNA^fMet^ (CAU), tRNA^Gly^ (GCC, UCC), tRNA^His^ (GUG), tRNA^Ile^ (CAU, GAU), tRNA^Lys^ (UUU), tRNA^Leu^ (UAA), tRNA^Asn^ (GUU), and tRNA^Val^ (GAC, UAC) (Supplementary File 9). Among them, tiRFs of tRNA^His^ (GUG) and tRNA^fMet^ (CAU) were found only once, whereas, tRNA^Lys^ (UUU) was the highest (72) encoding tiRF. The tiRFs of tRNA^Lys^ (UUU) was followed by tRNA^Ile^ (GAU) and tRNA^Ala^ (UGC), which were found to contain 51 and 52 putative tiRFs, respectively (Supplementary File 9).

### Machine Machine-learning approach showed GC% influences the tRNA number in the chloroplast genome

We grouped the chloroplast genomes of all the species according to their clade and conducted a comparative study. The analysis revealed that the average tRNA gene number in monocot (37.80%) plants is comparatively higher than that in other plants (Supplementary File 6). The protists showed the lowest (29.5%) average tRNA gene number, followed by algae (30.12%) (Supplementary File 6). A correlation analysis of the GC% with the tRNA number showed a positive correlation (r = 0.362) for the monocot clade (Figure 6). The chloroplast genomes of the species Isolepis setacea and Vitis romanetii were found to encode the highest number of tRNAs, that is, 52 each (Supplementary File 6). On an average, the chloroplast genomes were found to encode 36 tRNA genes per genome. A machine-learning approach was used to understand the role of the GC content and genome size in the tRNA number in the chloroplast genome. The boosting analysis revealed that the relative influence of the GC% was more than the genome size (Figure 7). A principal component analysis was conducted to see their association with different clades.

**Figure 6.**
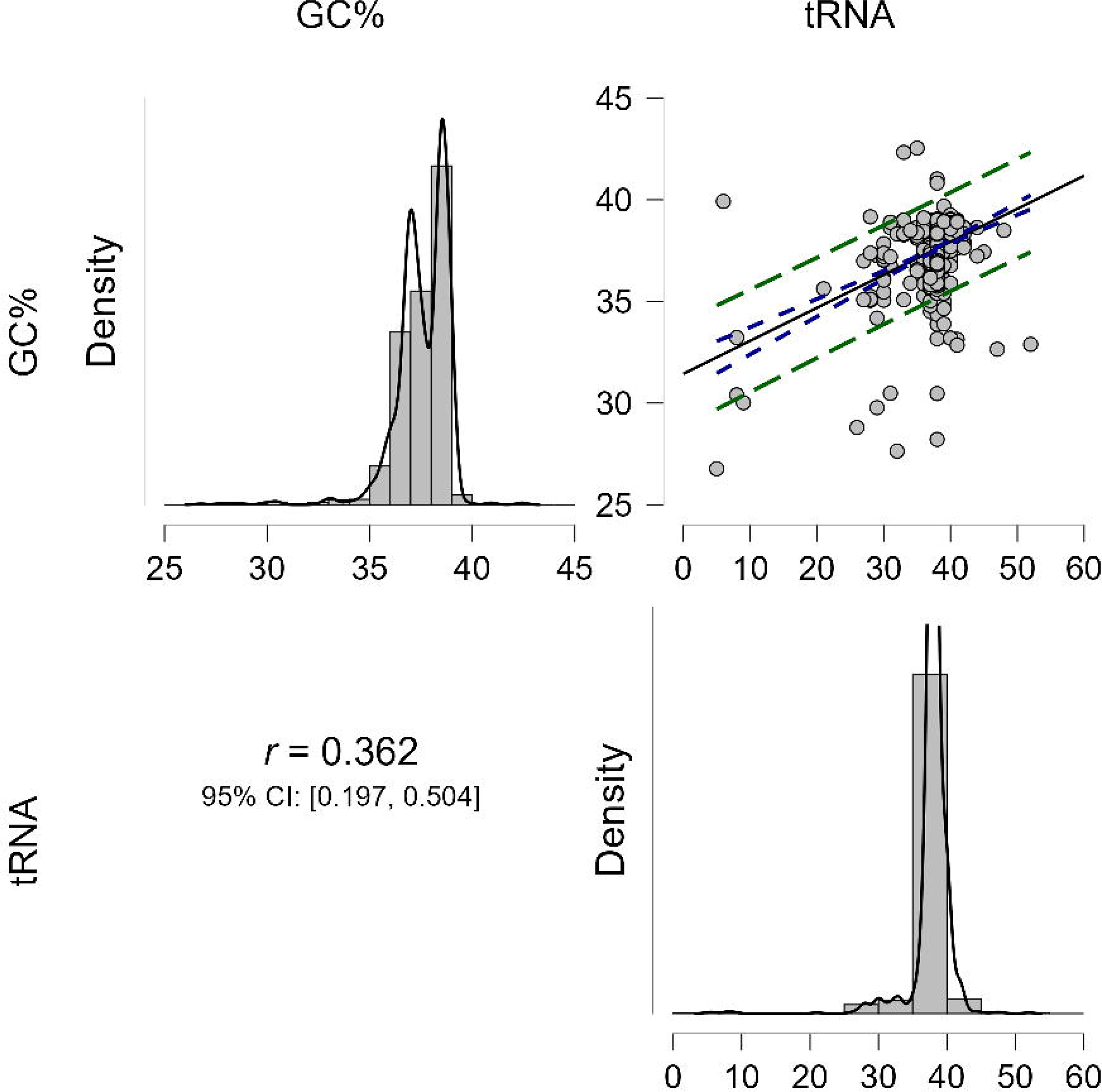
Correlation regression analysis (r = 0.362) of GC % and tRNA gene number in the chloroplast genome. Analysis showed that there was a slight positive correlation between the GC% and tRNA gene number in the chloroplast genome. The analysis was conducted at a p-value < 0.05. Correlation analysis was conducted using the JASP 0.16.1.0 version software.

**Figure 7.**
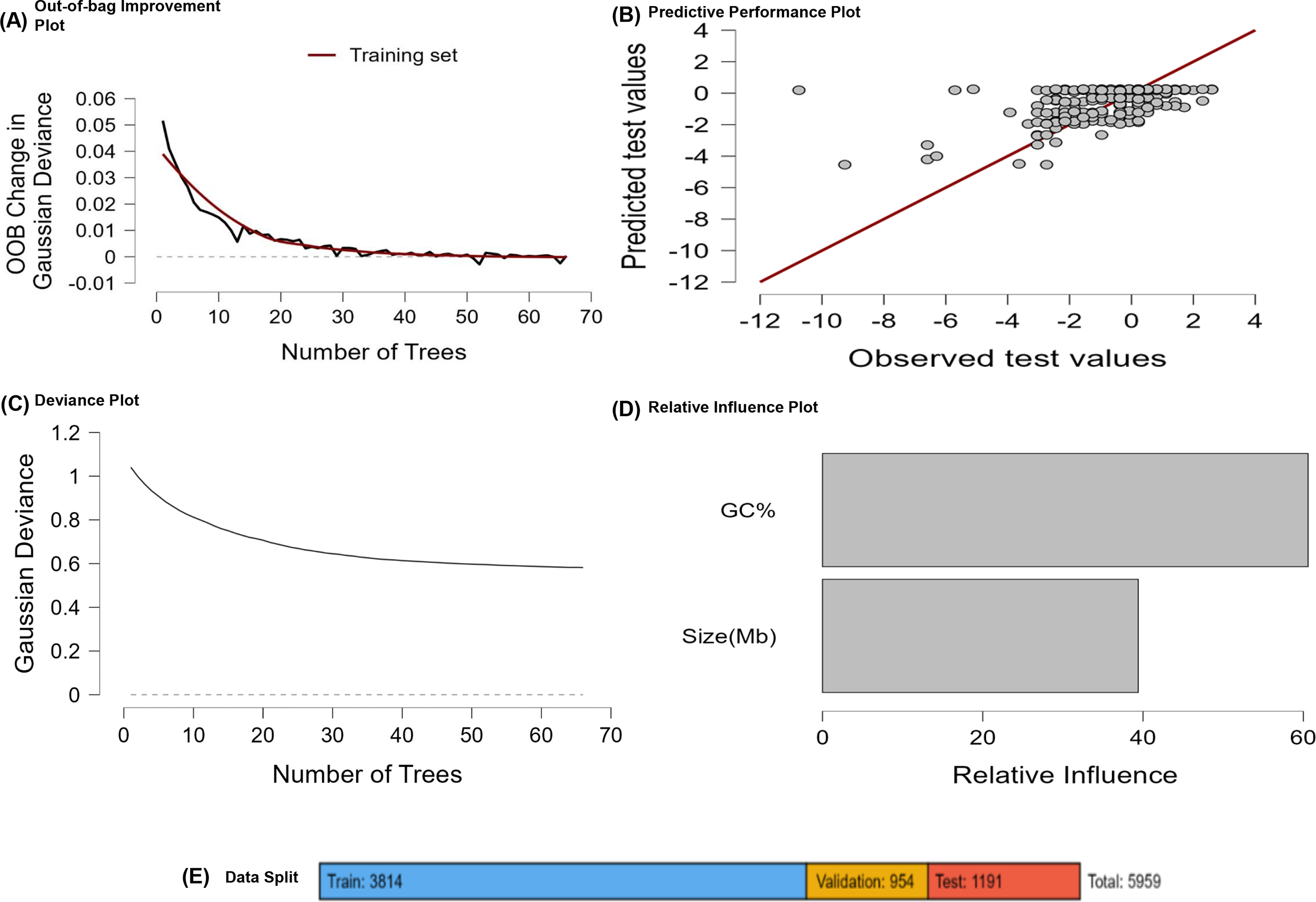
Machine-learning analysis of GC % content and genome size in the tRNA gene number in the chloroplast genome; the random forest approach was used to run the analysis. Analysis revealed that the GC% content had more influence toward the number of tRNA gene numbers than the genome size. In the study, from 5959 species, 3814 species were used as training sets, 954 for validation, and 1191 as test sets. All the analysis was conducted at p < 0.05.

### Chloroplast tRNAs Evolve from Multiple Common Ancestors

We conducted a phylogenetic analysis by considering the tRNA genes of the chloroplast genome. The phylogenetic analysis revealed clear and distinct phylogenetic clusters of tRNAs. The phylogenetic tree showed two major distinct clusters suggesting their origin from multiple common ancestors (Figure 8). In cluster I, anticodons GCU, GGA, UGA, GCC, UCC, CGU, CGA, GCC, CGA, CGU, UUC, UCU, CAU, UAA, CAA, GUA, UAG, UAU, UAUG, CAA, GCU, UCG, UCU, GAA, CUA, UAG, and GAG, grouped together, whereas, in cluster II, anticodons UUG, GUG, GCA, GAA, UUU, GUU, UGG, GGG, CCA, UGU, GGU, CAU, UAC, GCC, GUC, GAC, GAU, UUC, CGU, ACG, CCG, ACA, and UGC, grouped together (Figure 8). The anticodons GAA (tRNA^Phe^), CAU (tRNA^Met^), GCC (tRNA^Gly^), UUC (tRNA^Glu^), and CGU(tRNA^Thr^) were shared in both the clusters. The phylogenetic analysis of quadruplet anticodons revealed that quadruplet anticodon AUAA shares a phylogenetic relationship with UAUG anticodons, whereas, the UGGG and GUUA anticodons fall in a distinct cluster (Figure 9).

**Figure 8.**
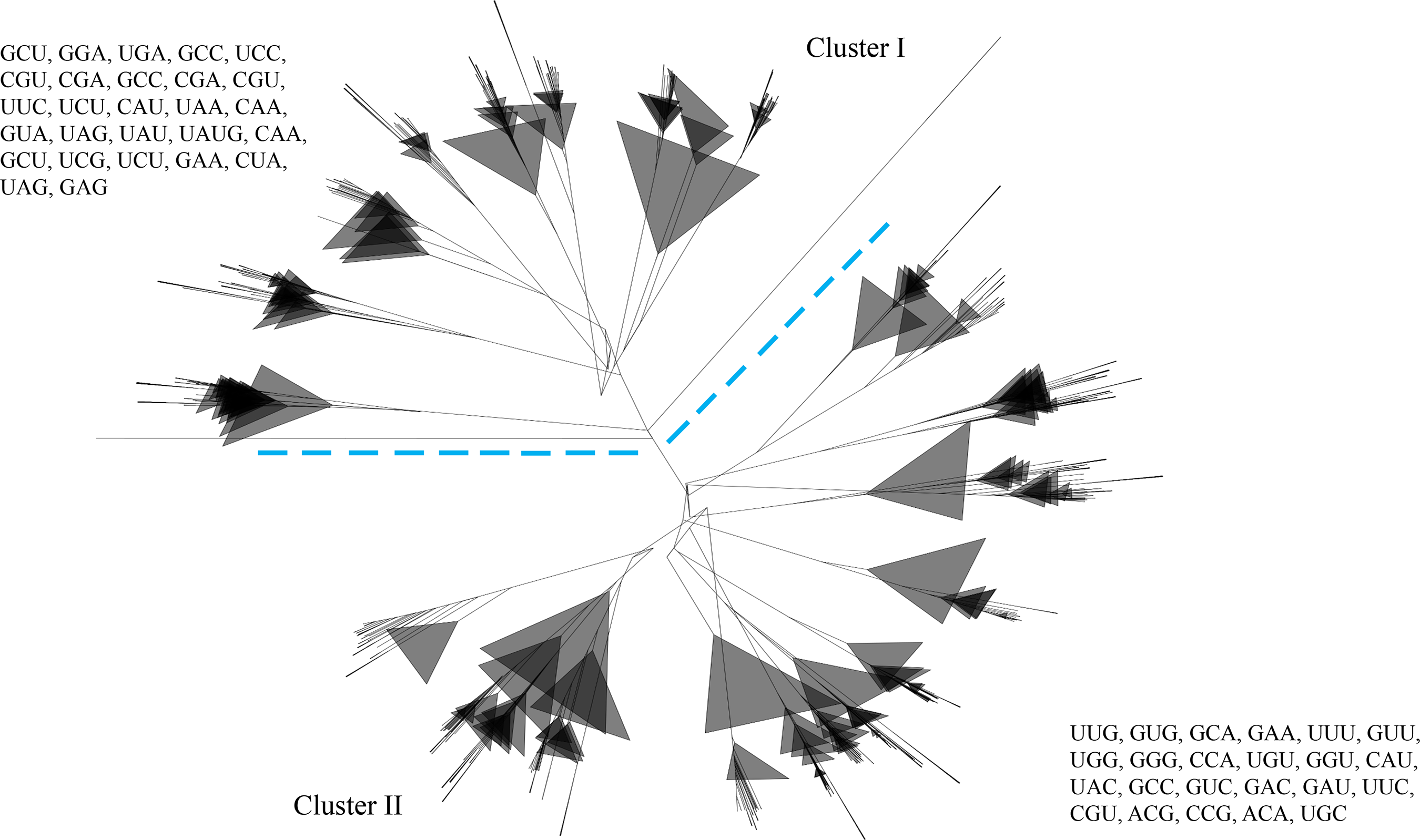
Phylogenetic tree of chloroplast tRNAs. The phylogenetic tree shows two distinct major clusters named cluster I and cluster II. The phylogenetic tree shows that chloroplast tRNAs have evolved from multiple common ancestors. In cluster I anticodons GCC, CGU, CGA, UCU, CAA, and UAG, are found in more than one group, and the anticodons GCC, GCU, UUC, CAU, and GAA are found in both the clusters, showing their evolution via duplication. The phylogenetic tree has been constructed using the neighbor-joining method, using the Clustal W program.

**Figure 9.**
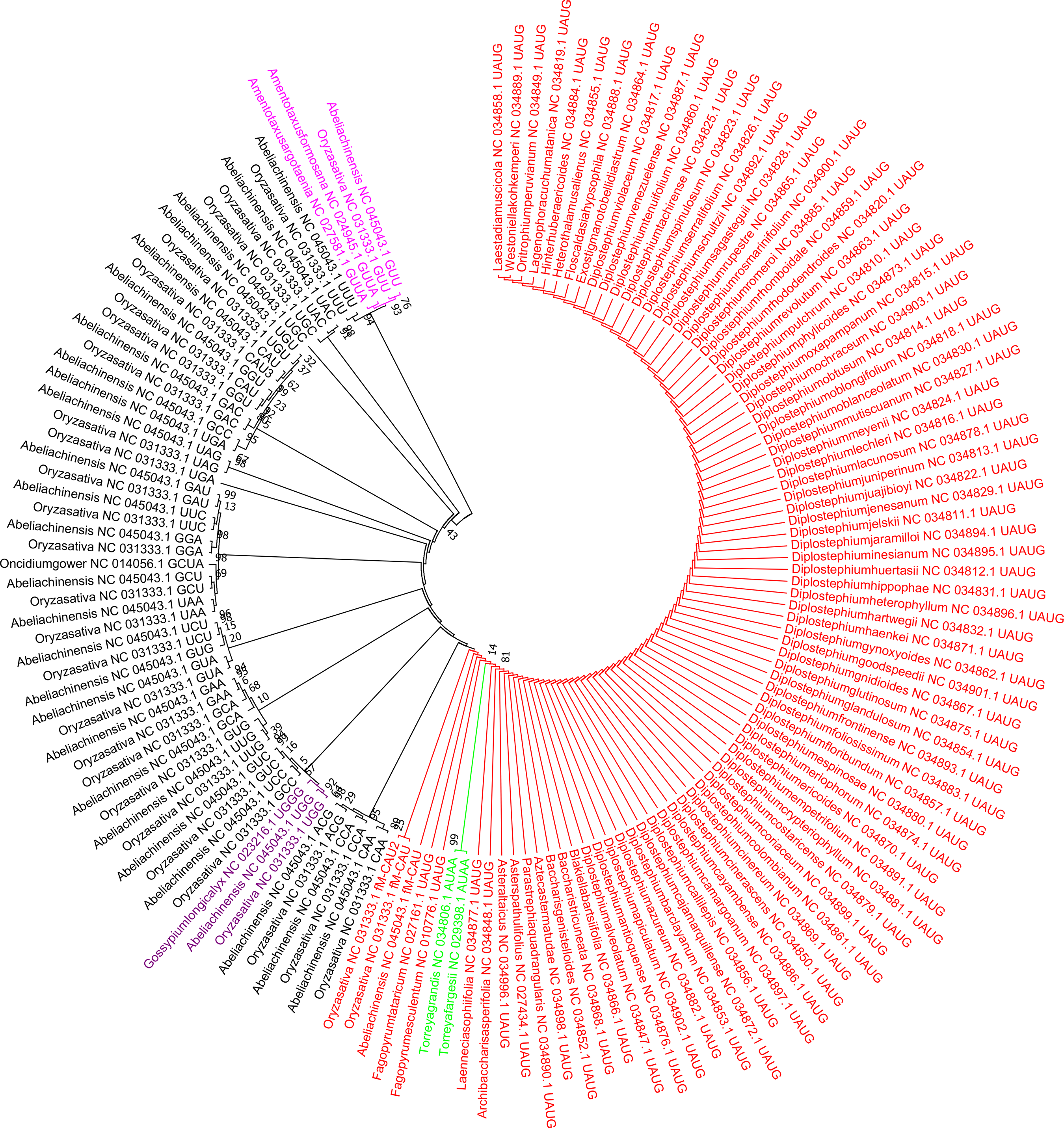
Phylogenetic tree of a putative quadruplet anticodon containing tRNAs with triplet codon– containing tRNAs. The phylogenetic grouping revealed that the quadruplet anticodons had evolved via addition of a nucleotide preceding the third nucleotide of the triplet anticodons. The evolutionary history was inferred by using the Maximum Likelihood method based on the Tamura-Nei model. The tree with the highest log likelihood (−1053.93) is shown. Initial tree(s) for the heuristic search were obtained automatically by applying the Neighbor-Join and BioNJ algorithms to a matrix of pair-wise distances, estimated by using the Maximum Composite Likelihood (MCL) approach, and then selecting the topology with the superior log likelihood value. The tree was drawn to scale, with branch lengths measured in the number of substitutions per site. The analysis involved 147 nucleotide sequences. All positions with less than 95% site coverage were eliminated, that is, fewer than 5% alignment gaps, missing data, and ambiguous bases, were allowed at any position. There were a total of 26 positions in the final dataset. Evolutionary analyses were conducted using the MEGA 7.

Genes undergo mutation, which is a common phenomenon. Although it was a common phenomenon in coding genes, non-coding genes also showed frequent mutation. Therefore, a transition/transversion bias study was conducted for the chloroplast tRNAs. The analysis revealed that transition predominates transversion (Supplementary Table 2). The transition/transversion bias was found to be the highest for tRNA^Asn^ (R = 13.71), whereas, tRNA^Ser^ (1.22) had the lowest bias (Supplementary Table 2). The transition/transversion bias of tRNA^Asn^ was followed by tRNA^Tyr^ (11.51) and tRNA^Trp^ (8.63). Although, tRNA^Arg^, tRNA^Leu^, and tRNA^Ser^ encoded six Isoacceptors, their transition/transversion bias was comparatively lower than that of others (Supplementary Table 2).

## Discussion

The chloroplast genome harbors several coding sequences and a few non-coding sequences including rRNA and tRNA. These genetic elements and their potential to translate codons make them semi-autonomous organelles of the plant cell. A detailed genomic analysis of the chloroplast tRNA reveals that it does not encode all the 64 anticodons required for the tRNAs. The tRNAs with anticodons ACU (tRNA^Ser^), CUG (tRNA^Gln^), GCG (tRNA^Arg^), CUC (tRNA^Glu^), CCC (tRNA^Gly^), and CGG (tRNA^Pro^), are absent in the chloroplast genome of the studied species. therefore, these anticodons can be classified as rare anticodons of the chloroplast genome. The ACU anticodon of tRNA^Ser^ and the GCG anticodon of tRNA^Arg^ are from the hexa-isoacceptor group, whereas, the CCC anticodon of tRNA^Gly^ and the CGG anticodon of tRNA^Pro^ are from the tetra-isoacceptor group. Therefore, a lack of these anticodons from their isoacceptor group does not make any difference in the genome as other isoacceptors are available for their use, to encode the codon. However, tRNA^Gln^ is encoded only by CUG and UUG anticodons, whereas, tRNA^Glu^ is encoded by the CUC and UUC anticodons. The lack of the CUG anticodon from tRNA^Gln^ and the CUC anticodon from tRNA^Glu^ in the chloroplast genome has left these tRNA isotypes with only one choice of anticodon (Table 1). The lack of the CUG anticodon in tRNA^Gln^ and the CUC anticodon in tRNA^Glu^, in the chloroplast genome, may be due to a strong selection pressure to establish UUG (tRNA^Gln^) and UUC (tRNA^Glu^) anticodons as the dominant anticodons. The tRNA anticodons followed by nucleotides CUx (x = any nucleotide) may have undergone a strong evolutionary pressure, and hence, anticodons CUA, CUU, CUG, and CUC, encode only 197, 7, 0, and 0 anticodons, respectively, in the chloroplast genome (Table 1). However, the CAU anticodon encoding tRNA^Met^ has been seen to have the highest percentage (5.47%) in the chloroplast genome (Supplementary File 1). The CAU anticodon of tRNA^Met^, of the nuclear encoded genome has also been found in the highest (5.03%) abundance [10], thus corroborating CAU, as the most abundant anticodon in the nuclear and chloroplast genomes. The anticodons CAU (tRNA^Met^), GUU (tRNA^Asn^), UGC (tRNA^Ala^), and ACG (tRNA^Arg^) have been found to encode more than 5% each of the total anticodons, suggesting the role of positive selection pressure in these anticodons (Supplementary File 1). However, at the isotype/isodecoder level, tRNA^Leu^ (10.27%) has been found to contain the highest percentage of anticodons followed by tRNA^Ile^ (9.93%) and tRNA^Arg^ (7.96%) (Supplementary File 1). A similar level of abundance has been found for tRNA^Leu^ (7.80%), for the nuclear encoded tRNA genes, reflecting a similarity in the anticodon abundance in the nuclear and chloroplast genomes [10]. However, an abundance of the nuclear-encoded anticodons tRNA^Leu^ is followed by tRNA^Ser^ (7.66%), tRNA^Gly^ (7.52%), and tRNA^Arg^ (7.28%) [10]. Although, tRNA^Leu^ is the highest encoding isotype/isodecoder in nuclear-(7.80%) and chloroplast (10.27%)-encoded genomes, there is a great difference in their percentage. The chloroplast-encoded CAU anticodon also encodes tRNA^Ile2^ (4.93%). The CAU anticodon for tRNA^fMet^ (0.33%) is also quite abundant in the chloroplast genome. The tRNA^fMet^ acts as an initiation anticodon in protein synthesis in mitochondria, bacteria, and chloroplasts and the presence of tRNA^fMet^ in the chloroplast genome is quite justified. However, only 709 tRNA^fMet^ genes were found during the analysis suggesting that tRNA^fMet^ is not a universal tRNA of the chloroplast genome. A majority percentage of the chloroplast genome does not encode tRNA^fMet^. A few of the chloroplast genomes encode the tRNAs for selenocysteine and pyrrolysine amino acid (Table 1). However, Zhao et al., (2021) has reported the absence of tRNA^Sec^ in gymnosperm plants [11]. The Sec amino acid specified by the UGA codon, requires the presence of the selenocysteine insertion sequence (SECIS) element, and the Pyl amino acid encoded by the UAG codon requires the pyrrolysine insertion sequence (PYLIS) [12]. The presence of tRNA for encoding Sec and Pyl reflects that the chloroplast genome may have SECIS and PYLIS in it.

It was also very peculiar to see the loss of tRNA genes in the chloroplast genome of heterotrophic and mycoparasitic plants. Our previous study reported the loss of several other genes in the chloroplast genome in mycoparasitic and heterotrophic plants [13]. Similar is true for the tRNA genes as well. In the absence of tRNA genes in the chloroplast genome, the cell most probably uses the tRNA genes from the nuclear-encoded genome. However, the loss of tRNA genes in the chloroplast genome seems independent of the nuclear genome. The parasitic and heterotrophic plants require less effort to complete their lifecycle, as they are completely dependent on their host. Hence, they do not need a lot of genes for their function, and hence, may be under constant pressure to eliminate genes. Therefore, these mycoparasitic and heterotrophic plants contain only 14 (CAU, CCA, GAA, GCA, GCU, GUA, GUC, GUG, GUU UCC, UCU, UUC, UUG, and UUU) anticodons in their chloroplast genome.

It is well known that the triplet genetic code is canonical and not universal. The genetic code can be expanded, where specific codons can be re-allocated to encode non-proteogenic amino acids. The tRNA genes undergo rapid changes to meet the translational demand of the cell [14]. Therefore, it is highly possible that tRNA can expand its anticodon nucleotide number. Our study helped us to discover the presence of quadruplet anticodons in the chloroplast genome of at least 91 plant species (Supplementary File 4). The quadruplet anticodons found in our study were UAUG, UGGG, AUAA, GCUA, and GUUA. Studies regarding the presence of functional quadruplet anticodons are reported in a few cases [15–22]. Anderson et al., (2004) reported the role of the quadruplet codon AGGA through changes in the tRNA anticodon loop to CUUCCUAAA in a suppressor tRNACUA [15]. The suppression of the amber tRNA led to the encoding of homoglutamine (hGln), using the AGGA codon [15]. They also reported that quadruplet codons CCCU or CUAG could be used to suppress the amber tRNA and allow the incorporation of unnatural amino acid into the protein in Escherichia coli[15]. Neumann et al., (2010), reported the encoding of unnatural amino acids through the evolution of the quadruplet anticodon in response to the amber codon tRNACUA[16]. Chloramphenicol resistance was achieved when tRNAUCUU^Ser2^ translated the AAGA codon and tRNAUCCU^Ser2^ translated the AGGA codon [16]. Niu et al., (2013) replaced tRNA^Pyl^CUA with the UCCU anticodon and generated tRNA^Pyl^UCCU, which recognized and suppressed the quadruplet codon AGGA [17]. This provided a qualitative notion for the suppression of the quadruplet codon through tRNAUCCU [17]. Most specifically, the presence of the quadruplet anticodon was associated with suppression of the amber tRNA and incorporation of the unnatural amino acid into the protein chain. The tRNAGCUA contained an additional G nucleotide prior to the tRNACUA anticodon, suggesting its role in suppression of the amber codon. In the tRNA^Asn^GUUA anticodon, most probably, nucleotide A was incorporated after the GUU anticodon, as the tRNA with the GUU anticodon was grouped with the GUUA anticodon in the phylogenetic tree (Figure 9). Similarly, in the UGGG anticodon, the G nucleotide got incorporated in the UGG anticodon, as they grouped with the UGG anticodon (Figure 9). The GCUA anticodon was grouped with GCU anticodon suggesting that the A nucleotide was incorporated at the fourth position of the GCU anticodon, which gave rise to the GCUA anticodon (Figure 9). However, no such clue was found in the case of the UAUG and AUAA anticodons. Considering, the incorporation of the additional nucleotide at the fourth position, we could speculate that the G nucleotide was most probably incorporated in the UAU anticodon, and gave rise to the UAUG anticodon. Similarly, the A nucleotide was incorporated at the 4^th^ position of the AUA anticodon to give rise to the AUAA anticodon. Although, we found only five putative quadruplet anticodons, the genome could accommodate at least 256 quadruplet anticodons/codons in the cell (Table 2). We also found the presence of tRNAs, with only duplet anticodon, where one nucleotide was possibly deleted from the anticodon (Supplementary File 5). At least 13 species resulted that contained duplet anticodons in the tRNA of the chloroplast genome (Supplementary File 5).

The chloroplast encoding tRNAs were also found to encode the group I introns. These group I introns were conserved in their respective isotype/isodecoder groups (Table 2). From a total of 20 isotypes, 12 of them were found to encode the group I introns (Table 2). However, the group I intron of one isotype was not conserved with the intron of another isotype, reflecting the isotype-based conservation of the group I intron, in the tRNA.

It is well-reported that group I introns are found in tRNAs, bacteria, lower eukaryotes, and higher plants [23–25]. Some of the group I intron encode homing endonucleases catalyze intron mobility, thus facilitating the movement of the intron from one location to another and from one organism to another [24]. However, the incorporation of the group I intron in the tRNA gene is isotype-specific, as only 12 isotypes have been found to encode the intron, while eight isotypes do not have any intron in their tRNAs (Table 2). From the eight isotypes, tRNA^His^, tRNA^Gln^, tRNA^Asp^, tRNA^Asn^, and tRNA^Arg^ belong to the polar group, whereas, tRNA^Trp^, tRNA^Pro^, and tRNA^Val^ belong to the non-polar group. This shows that the presence of the type I intron tends to be more toward the tRNA that encodes polar amino acids. Furthermore, it is seen that the chloroplast genome also encodes the putative spacer tRNAs (Figure 2). It is reported that E. coli contains a spacer tRNA (tRNA^Ala^ and tRNA^Ile^) that is present in the spacer region of the 16S and 23S rRNA [26]. The tRNAs, tRNA^Ala^, and tRNA^Ile^, have also been found in the spacer region of 16S and 23S rRNA suggesting the presence of a spacer tRNA in the chloroplast genome. Although, in a majority of cases, tRNA^Ala^ and tRNA^Ile^ are the predominant spacer tRNAs; tRNA^Glu^ can be the third most possible spacer tRNA of the chloroplast genome.

Analysis also revealed the presence of tRNA fragments (tRFs) in the chloroplast genome. We found at least 55 tRFs that belonged to ten tRNA isotypes (Supplementary File 8). These tRFs were putatively derived from the tRNA precursors or from the cleavage of mature tRNAs [27]. The tRFs were reported to control gene expression, translation control, transposon control, ncRNA, and DNA damage response [8, 27–29]. Although, we found ten different chloroplast-derived tRFs, the majority of them belonged to tRNA^Glu^ and tRNA^Gly^ (Supplementary File 8). Among them are the, tRNA^Glu^are tRF-1 type, tRNA^Gly^are tRF-5ˈ-type, and tRNA^His^, tRNA^Thr^, and tRNA^Val^are tRF-3ˈ type (Supplementary File 8). Furthermore, we also noted the presence of a few putative tRNA-derived, stress-induced RNA (tiRNAs) fragments (tiRFs) in the chloroplast genome. The majority of the tiRFs were from tRNA^Lys^ (UUU). For the first time, tiRFs were reported in the human fetus hepatic tissue and osteosarcoma cells [30, 31]. These tiRFs could be generated in the cell under different stress conditions via cleavage of mature tRNAs [30]. However, their presence as independent nucleotide fragments in the annotated genome sequence reflected their independent presence in the genome. Although, the cleavage of tRNAs to tiRFs was brought about by the enzyme angiogenin (an RNase superfamily) [31] in the human cell; its counterpart in plants needs to be identified to understand its detailed functions. The 5ˈ-tiRNA^Ala^ and tiRNA^Cys^ were reported to inhibit translation in rabbit reticulocytes [31] suggesting their inhibitory role in protein translation.

This study also found the presence of a putative novel tRNA structure encoded by the chloroplast genome (Figure 4). The tRNA^Gly^ (UCC) was found to contain a long nucleotide sequence between the D-arm and anticodon arm in several species. This long arm could be most probably be an intron that might have incorporated in between these two arms. The chloroplast tRNAs which had lost the pseudouridine loop (Ψ) seemed to be metazoan mitochondrial-specific (Figure 5). The loss of the Ψ-loop in tRNA was first reported in the 1970s [32–34]. Previous studies also reported loss of the Ψ-arm and loop in nematode mitochondrial tRNA [34]. However, in the nematode mitochondrial tRNA, the Ψ-arm and loop were present in the tRNA^Ser^(GCU), whereas, it had lost the Ψ-arm and loop in tRNA^Ser^ (GCU and GGA) in the chloroplast genome (Figure 5). The elongation factor (EF) Tu combined with GTP to form a complex that delivered the amino acyl tRNA to the ribosome A site through binding of the acceptor arm and Ψ-arm [35]. In the absence of the Ψ-arm and loop in the tRNAs, it might be using some alternative binding mode for EF-Tu [36, 37]. In the case of Caenorhabditis elegans mitochondrial EF-Tu, it has around 60 amino acid extensions at the C-terminal end that might be playing important role in binding tRNAs that lack the Ψ-arm [38, 39]. This also suggested that the mitochondrial ribosomal protein might have alternate binding sites for the truncated tRNA. Furthermore, the presence of the metazoan, mitochondria-specific, truncated tRNA in the chloroplast genome suggested that these tRNA genes might be shared by sub-cellular organelle chloroplast and mitochondria.

Evolutionary analysis revealed, chloroplast tRNAs are derived from multiple common ancestors (Figure 8). The phylogenetic tree of the chloroplast tRNA shows two distinct clusters, which reflect their evolution from multiple common ancestors. In cluster I, anticodons GCC, CGU, CGA, UCU, CAA, and UAG are seen to make more than one group, whereas, none of the anticodons from cluster II are found to make more than one group (Figure 8). The anticodons GCC, GCU, UUC, CAU, and GAA are also found in both the clusters (Figure 8). This suggests that tRNAs with anticodons GCC, CGU, CGA, UCU, CAA, and UAG, of cluster I, may have undergone vivid duplication and produced more than one anticodon group.

## Conclusions

Chloroplast is a semiautonomous organelle of the plant and protist kingdom with a great potential to encode its own genome and protein translation machinery. The important tRNA molecules require for protein translation process is well documented. Chloroplast genome encode putative duplet, triplet, and quadruplet anticodons suggesting their role in recognition of duplet, triplet, and quadruplet codons in the mRNA. Mycoparasitic plants has lost their chloroplast genome to a large extent thereby losing several chloroplast encoded tRNA genes. Further, several of the chloroplast encoded tRNA genes were found to encode introns and the presence of intron in chloroplast genome suggest the presence of introns in the gene of their prokaryotic ancestor cyanobacteria. Further, the chloroplast genome is very selective and encoded only a few Isoacceptor abundantly while GCG, CUG, CUC, CCC, CGG, and ACU anticodons were found to be the rarest form of anticodons in the chloroplast genome. It is important to understand why chloroplast genome do not encode tRNA with such anticodons.

## Materials and Methods

All the chloroplast genomes were downloaded from the National Center for Biotechnology Information (NCBI) database. In total, 5959 chloroplast genomes were used in this study. The downloaded chloroplast genomes were subjected to tRNA annotation. tRNA annotation was conducted using tRNAscan-SE 2.0, Aragorn and the GeSeq-Annotation of the organellar genomes [40–42]. The Linux-based approach was used to annotate the chloroplast tRNA for tRNAscan-SE 2.0 and Aragorn. In the GenSeq-annotation of the organellar genome, the chloroplast genome files were uploaded with the following parameters; sequence source: Plastid; annotation option: Annotate plastid inverted repeats; blat search: Default; annotate: CDS, tRNA, and rRNA; and third party tRNA annotator: Aragorn v1.2.38, tRNAscan-SE v2.0.7. All the tRNA sequences generated from these three annotation pipelines were corroborated and used for further analysis. All the data obtained from tRNAscan-SE and Aragorn were further processed in an excel worksheet. The Organellar Genome Draw (OGDRAW) was used to draw the organellar genome map of the chloroplast genome [43]. The Genbank file was used to draw the chloroplast genome map in OGDRAW [43].

### Multiple sequence alignment

The intron sequences retrieved from the chloroplast tRNA were aligned to find the possible conserved structure. Multiple sequence alignment was conducted using the Multalin software (http://multalin.toulouse.inra.fr/multalin/) that uses hierarchical clustering [44]. Default parameters were used to construct the alignment.

### Machine Learning Approach and Statistical Analysis

A machine learning approach was used to understand the role of the genome size and GC% content in the number of tRNA genes in the chloroplast genome. The random forest regression approach was used for this purpose. The following parameters were used in the random forest analysis: target tRNA gene number, predictor’s genome size, and GC% content; Plots: data split, out-of-bag error, predictive performance, mean decrease in accuracy, and total increase in node purity; tables: evaluation matrix; data split preference: sample 20% of all data; training and validation of data: 20% validation data. The training parameters were as follows, training data used per tree: 50%; predictor per split: auto; and max tree: 100%. The machine-learning approach was studied using the JASP software version 0.16.1.0 [45]. The correlation plot for GC% content and tRNA was also conducted using the JASP 0.16.1.0 software. The following parameters were used for the correlation analysis, sample correlation coefficient: Pearson’s r and confidence interval: 95% (p < 0.05)[45].

### Phylogenetic tree

The tRNA sequences of the chloroplast genomes were taken to construct the phylogenetic tree. The phylogenetic tree was constructed using the Clustalw program in a Linux-based environment. A neighbor joining tree was constructed with 100 bootstrap replicates. The resulting file was saved in nwk file format and later uploaded in the iTOL Interactive Tree of Life, to view the tree [46]. The phylogenetic tree of the tRNA quadruplet anticodons, with other anticodons, was constructed using the MEGA software version 7[47]. Prior to the construction of the phylogenetic tree, the tRNA sequences were subjected to multiple sequence alignments. Multiple sequence alignments were conducted using the MUSCLE software[48]. The resulting clustal file was converted to the MEGA file format (aln) using the MEGA 7 software[47]. The converted file was subjected to construct the phylogenetic tree in the MEGA 7 software, using the maximum-likelihood approach. The Tamura-Nei model, with a 500-bootstrap replicate, was used for this analysis. The phylogenetic tree of the tRNA introns was also constructed using the MEGA 7 software with the same statistical parameters [47].

## Acknowledgements

Acknowledgement

N/A

## Statement and Declarations

## Data availability

All the data used during this study was taken from National Center for Biotechnology Information database and all the data are available in the public domain. Also, the accession numbers are provided in the supplementary files.

## Supplementary Materials

**Supplementary File 1.** Percentage of anticodons in the chloroplast genome.

**Supplementary File 2.** Name of the species with rare anticodons in their tRNA gene.

**Supplementary File 3.** Name of the species encoding UCA anticodon for tRNA selenocysteine.

**Supplementary File 4.** Name and accession number of the species encoding putative quadruplet anticodons in the chloroplast genome.

**Supplementary File 5.** Accession number of the species encoding putative duplet anticodons in the chloroplast tRNA.

**Supplementary File 6.** Genomic details of chloroplast genome of different species. Supplementary File 7. List of species those do not contain the spacer tRNA. Supplementary file 8. The list of tRNA fragments found in the chloroplast genome. Supplementary File 9. Putative tiRNAs of chloroplast genome.

**Supplementary Table 1.** Quadruplet anticodon/codon table. There are 256 possibilities to encode an amino acid via quadruplet anticodon/codon. It can accommodate maximum of the amino acids available in the proteome to its protein translation machinery.

**Supplementary Table 2**

Transition and transversion bias (MCL) of different tRNA genes of the chloroplast genome. Each entry shows the probability of substitution (r) from one base (row) to another base (column). For simplicity, the sum of r-values is made equal to 100. Rates of different transitional substitutions are shown in bold and those of transversionsal substitutions are shown in italics. The transition/transversion rate ratios are k1 indicates purines and k2 indicates pyrimidines. The overall transition/transversion bias is mentioned as R where R = [A*G*k1 + T*C*k2]/[(A+G)*(T+C)]. All positions with less than 95% site coverage were eliminated. That is, fewer than 5% alignment gaps, missing data, and ambiguous bases were allowed at any position. The evolutionary analyses were conducted in MEGA7.

**Supplementary Figure 1.** Grouping of conserved type II introns found in tRNA of the chloroplast genome

