## Supplementary material for "Anticodon Table of the Chloroplast Genome and Identification of Putative Quadruplet Anticodons in Chloroplast tRNAs": Supplementary Figure 1.pptx

### Slide 1
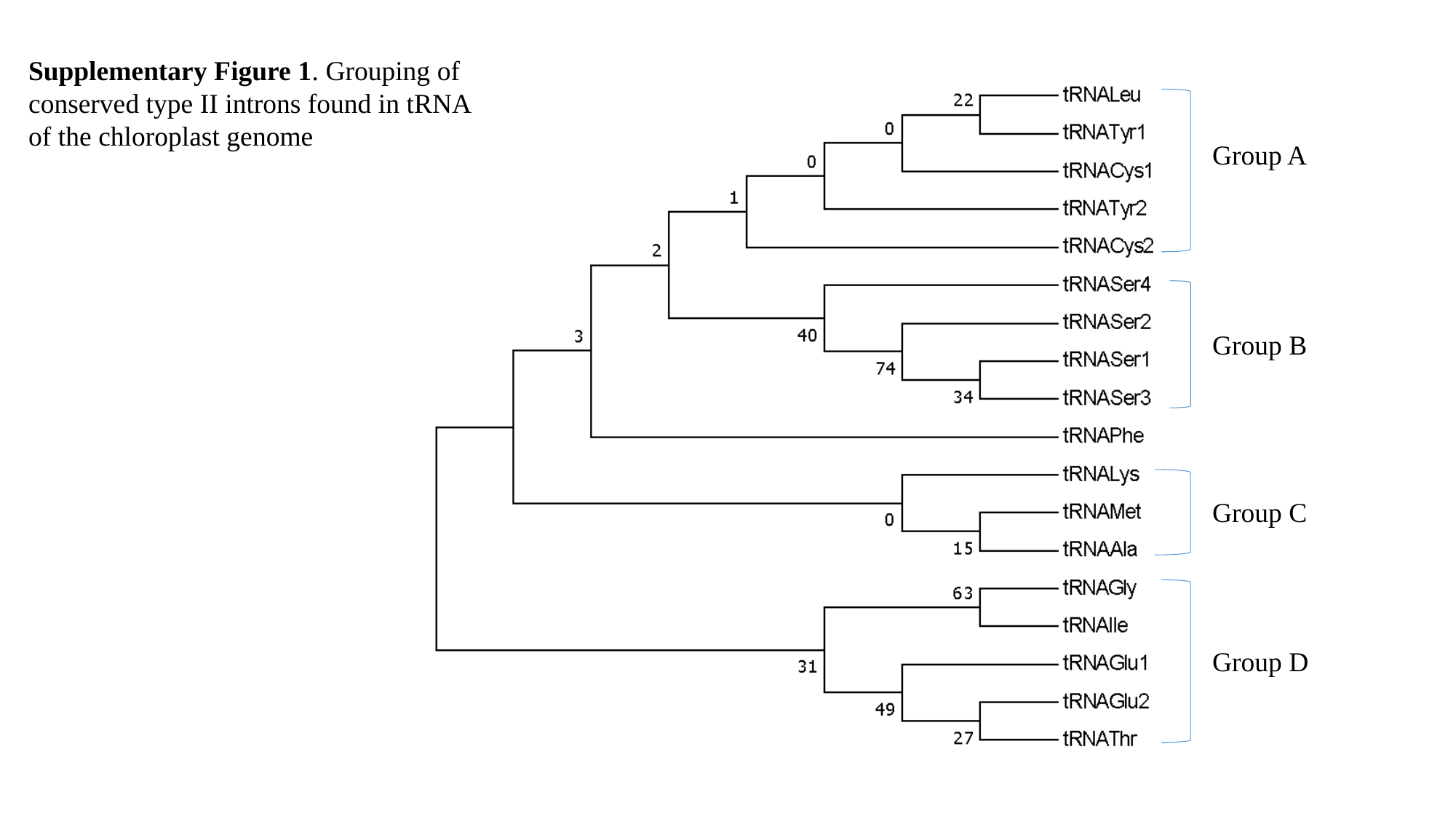

Supplementary Figure 1. Grouping of conserved type II introns found in tRNA of the chloroplast genome
Group A
Group B
Group C
Group D
