## Supplementary material for "Anticodon Table of the Chloroplast Genome and Identification of Putative Quadruplet Anticodons in Chloroplast tRNAs": Supplementary Table 1.docx

**Supplementary Table 1.** Quadruplet anticodon/codon table. There are 256 possibilities to encode an amino acid via quadruplet anticodon/codon. It can accommodate maximum of the amino acids available in the proteome to its protein translation machinery.

| UUUU | UGUU | UCUU | UAUU | GUUU | GGUU | GCUU | GAUU | CUUU | CGUU | CCUU | CAUU | AUUU | AGUU | ACUU | AAUU |
| --- | --- | --- | --- | --- | --- | --- | --- | --- | --- | --- | --- | --- | --- | --- | --- |
| UUUG | UGUG | UCUG | UAUG | GUUG | GGUG | GCUG | GAUG | CUUG | CGUG | CCUG | CAUG | AUUG | AGUG | ACUG | AAUG |
| UUUC | UGUC | UCUC | UAUC | GUUC | GGUC | GCUC | GAUC | CUUC | CGUC | CCUC | CAUC | AUUC | AGUC | ACUC | AAUC |
| UUUA | UGUA | UCUA | UAUA | GUUA | GGUA | GCUA | GAUA | CUUA | CGUA | CCUA | CAUA | AUUA | AGUA | ACUA | AAUA |
| UUGU | UGGU | UCGU | UAGU | GUGU | GGGU | GCGU | GAGU | CUGU | CGGU | CCGU | CAGU | AUGU | AGGU | ACGU | AAGU |
| UUGG | UGGG | UCGG | UAGG | GUGG | GGGG | GCGG | GAGG | CUGG | CGGG | CCGG | CAGG | AUGG | AGGG | ACGG | AAGG |
| UUGC | UGGC | UCGC | UAGC | GUGC | GGGC | GCGC | GAGC | CUGC | CGGC | CCGC | CAGC | AUGC | AGGC | ACGC | AAGC |
| UUGA | UGGA | UCGA | UAGA | GUGA | GGGA | GCGA | GAGA | CUGA | CGGA | CCGA | CAGA | AUGA | AGGA | ACGA | AAGA |
| UUCU | UGCU | UCCU | UACU | GUCU | GGCU | GCCU | GACU | CUCU | CGCU | CCCU | CACU | AUCU | AGCU | ACCU | AACU |
| UUCG | UGCG | UCCG | UACG | GUCG | GGCG | GCCG | GACG | CUCG | CGCG | CCCG | CACG | AUCG | AGCG | ACCG | AACG |
| UUCC | UGCC | UCCC | UACC | GUCC | GGCC | GCCC | GACC | CUCC | CGCC | CCCC | CACC | AUCC | AGCC | ACCC | AACC |
| UUCA | UGCA | UCCA | UACA | GUCA | GGCA | GCCA | GACA | CUCA | CGCA | CCCA | CACA | AUCA | AGCA | ACCA | AACA |
| UUAU | UGAU | UCAU | UAAU | GUAU | GGAU | GCAU | GAAU | CUAU | CGAU | CCAU | CAAU | AUAU | AGAU | ACAU | AAAU |
| UUAG | UGAG | UCAG | UAAG | GUAG | GGAG | GCAG | GAAG | CUAG | CGAG | CCAG | CAAG | AUAG | AGAG | ACAG | AAAG |
| UUAC | UGAC | UCAC | UAAC | GUAC | GGAC | GCAC | GAAC | CUAC | CGAC | CCAC | CAAC | AUAC | AGAC | ACAC | AAAC |
| UUAA | UGAA | UCAA | UAAA | GUAA | GGAA | GCAA | GAAA | CUAA | CGAA | CCAA | CAAA | AUAA | AGAA | ACAA | AAAA |
