## Supplementary material for "Anticodon Table of the Chloroplast Genome and Identification of Putative Quadruplet Anticodons in Chloroplast tRNAs": Supplementary Table 2.docx

|  | A | T | C | G | A | T | C | G | A | T | C | G |
| --- | --- | --- | --- | --- | --- | --- | --- | --- | --- | --- | --- | --- |
|  | **Asn** (k1=19.371, k2=38.362, R=13.71) | | | | **Asp** (k1=3.42, k2=13.46, R=3.961) | | | | **Met** (k1=5.16, k2=2.55, R=1.94) | | | |
| A | - | 0.9 | 0.68 | **20.98** | - | 2.04 | 2.69 | **11.08** | - | 4.43 | 3.94 | **21.62** |
| T | 0.61 | - | **25.99** | 1.06 | 1.72 | - | **36.22** | 3.24 | 4.45 | - | **10.05** | 4.19 |
| C | 0.61 | **34.59** | - | 1.06 | 1.72 | **27.44** | - | 3.24 | 4.45 | **11.31** | - | 4.19 |
| G | **11.94** | 0.9 | 0.68 | - | **5.89** | 2.04 | 2.69 | - | **22.99** | 4.43 | 3.94 | - |
|  | **Gln** (k1=7.38, k2=11.46, R=4.45) | | | | **Gly** (k1=4.23, k2=7.62, R=2.82) | | | | **Phe** (k=9.00, k2=20.64, R=7.07) | | | |
| A | - | 2.15 | 2.23 | **22.19** | - | 2.87 | 2.97 | **17.35** | - | 1.46 | 1.37 | **15.54** |
| T | 1.36 | - | **25.62** | 3.01 | 2.93 | - | **22.64** | 4.09 | 1.52 | - | **28.38** | 1.73 |
| C | 1.36 | **24.64** | - | 3.01 | 2.93 | **21.86** | - | 4.09 | 1.52 | **30.22** | - | 1.73 |
| G | **10.05** | 2.15 | 2.23 | - | **12.43** | 2.87 | 2.97 | - | **13.69** | 1.46 | 1.37 | - |
|  | **His** (k1=5.72, k2=9.34, R=3.7) | | | | **Ile** (k1=2.49, k2=2.79, R=1.31) | | | | **Pro** (k1=3.61, k2=6.73, R=2.5) | | | |
| A | - | 2.44 | 2.86 | **18.15** | - | 5.36 | 5.05 | **15.4** | - | 4.04 | 2.89 | **14.76** |
| T | 1.98 | - | **26.75** | 3.17 | 4.97 | - | **14.11** | 6.17 | 2.95 | - | **19.46** | 4.08 |
| C | 1.98 | **22.83** | - | 3.17 | 4.97 | **14.96** | - | 6.17 | 2.95 | **27.2** | - | 4.08 |
| G | **11.35** | 2.44 | 2.86 | - | **12.42** | 5.36 | 5.05 | - | **10.66** | 4.04 | 2.89 | - |
|  | **Trp** (k1=23.5, k2=12.46 R=8.63) | | | | **Tyr** (k1=0.791, k2=50.62, R=11.51) | | | | **Val** (k1=1.85, k2=3.79, R=1.42) | | | |
| A | - | 1.39 | 1.39 | **25.64** | - | 0.88 | 0.91 | **0.88** | - | 5.72 | 4.79 | **9.46** |
| T | 1.26 | - | **17.32** | 1.09 | 0.78 | - | **46.22** | 1.23 | 5.03 | - | **18.2** | 5.1 |
| C | 1.26 | **17.28** | - | 1.09 | 0.78 | **44.73** | - | 1.23 | 5.03 | **21.72** | - | 5.1 |
| G | **29.52** | 1.39 | 1.39 | - | **0.56** | 0.88 | 0.91 | - | **9.34** | 5.72 | 4.79 | - |
|  | **Arg** (k1=2.6, k2=4.09, R=1.64) | | | | **Leu** (k1=5.94, k2=0.59, R=2.08) | | | | **Thr** (k1=2.39, k2=18.69, R=5.39) | | | |
| A | - | 4.69 | 4.62 | **13.76** | - | 3.91 | 3.64 | **32.45** | - | 2.18 | 1.83 | **4.78** |
| T | 4.22 | - | **18.73** | 5.28 | 4.67 | - | **2.16** | 5.46 | 1.89 | - | **34.24** | 1.99 |
| C | 4.22 | **18.91** | - | 5.28 | 4.67 | **2.32** | - | 5.46 | 1.89 | **40.67** | - | 1.99 |
| G | **11** | 4.69 | 4.62 | - | **27.72** | 3.91 | 3.64 | - | **4.53** | 2.18 | 1.83 | - |
|  | **Ala** (k1=3.35, k2=6.78, R=2.63) | | | | **Lys** (k1=, k2=, R=4.683) | | | | **Ser** (k1=2.23, k2=2.69, R=1.22) | | | |
| A | - | 3.91 | 3.53 | **13.36** | - | 2.19 | 2.38 | **20.32** | - | 6.03 | 5.13 | **13.9** |
| T | 2.49 | - | **23.93** | 3.99 | 1.19 | - | **23.68** | 2.32 | 5.02 | - | **13.83** | 6.21 |
| C | 2.49 | **26.55** | - | 3.99 | 1.19 | **21.78** | - | 2.32 | 5.02 | **16.27** | - | 6.21 |
| G | **8.34** | 3.91 | 3.53 | - | **16.62** | 2.78 | 2.38 | - | **11.24** | 6.03 | 5.13 | - |
|  | **Cys** (k1=9.97, k2=0.26, R=2.74) | | | | **Glu** (k1=16.76, k2=3.89, R=4.24) | | | |  |  |  |  |
| A | - | 3.05 | 3.24 | **43.92** | - | 2.15 | 2.19 | **47.03** |  |  |  |  |
| T | 2.76 | - | **0.85** | 4.4 | 1.16 | - | **8.5** | 2.81 |  |  |  |  |
| C | 2.76 | **0.8** | - | 4.4 | 1.16 | **8.35** | - | 2.81 |  |  |  |  |
| G | **27.25** | 3.05 | 3.24 | - | **19.52** | 2.15 | 2.19 | - |  |  |  |  |

**Supplementary Table 2**

Transition and transversion bias (MCL) of different tRNA genes of the chloroplast genome. Each entry shows the probability of substitution (r) from one base (row) to another base (column). For simplicity, the sum of r values is made equal to 100. Rates of different transitional substitutions are shown in bold and those of transversionsal substitutions are shown in italics. The transition/transversion rate ratios are k1 indicates purines and k2 indicates pyrimidines. The overall transition/transversion bias is mentioned as R where R = [A*G*k1 + T*C*k2]/[(A+G)*(T+C)]. All positions with less than 95% site coverage were eliminated. That is, fewer than 5% alignment gaps, missing data, and ambiguous bases were allowed at any position. The evolutionary analyses were conducted in MEGA7.
